# Partial Bypass of Frataxin Deficiency by ISCU M141I Restores Cytosolic and Nuclear Fe–S Cluster Assembly

**DOI:** 10.1101/2025.09.03.673074

**Authors:** Valentine Mosbach, Nunziata Maio, Nadège Diedhiou, Adèle Hennick, Laure Dall’Agnol, Laurence Reutenauer, Lénaïc Marczak, Marie-Christine Birling, Aurélie Eisenmann, Alain Martelli, Puccio H Hélène

## Abstract

Iron-sulfur (Fe-S) clusters are essential cofactors required for the activity of numerous proteins involved in fundamental cellular processes, including DNA replication, metabolism and mitochondrial respiration. In eukaryotes, Fe-S cluster biogenesis is initiated in mitochondria by the ISC machinery, which assembles iron and sulfur, delivered by a cysteine desulfurase, onto the scaffold protein ISCU. Frataxin (FXN), a key regulator of this pathway, enhances Fe-S production by accelerating persulfide transfer to ISCU. FXN is essential in eukaryotes, and its loss results in “petite” phenotype in yeast, senescence in dividing mammalian cells and embryonic lethality in mice. Interestingly, in yeast, a methionine to isoleucine substitution at position 141 of the scaffold protein Isu1 can bypass the requirement of FXN. To test whether this bypass mechanism is conserved in mammals, we introduced the equivalent M141I substitution into the endogenous Iscu gene in murine fibroblasts carrying a conditional Fxn allele using CRISPR-Cas9. We show that the ISCU M141I variant enables cell survival in the absence of FXN, preventing cell cycle arrest and decreasing baseline DNA damage. However, these FXN-null survivor clones exhibit slower proliferation, persistent mitochondrial dysfunction and defective mitochondrial Fe-S cluster proteins. In contrast, nuclear and cytosolic Fe-S proteins are preserved, as is cellular iron homeostasis. Importantly, the ISCU M141I variant delays, but does not fully rescue, embryonic lethality in Fxn-deficient mice. Altogether, our results reveal a previously unrecognized compartment-specific rescue of Fe-S cluster dependent processes by the ISCU M141I variant in mammalian cells, raising for the first time the possibility of compartmental regulation of Fe-S cluster biogenesis.

## Introduction

Frataxin (FXN) is a highly conserved mitochondrial protein found across all domains of life, drawing significant attention due to its involvement in Friedreich’s ataxia (FA), a severe autosomal recessive neurodegenerative disorder^1,2^. FA is primarily caused by a GAA trinucleotide repeat expansion within the first intron of the *FXN* gene, leading to a marked reduction in FXN protein expression in patients^3^. Although FXN was initially characterized as an iron-binding protein and thereby was suggested to be involved in multiple aspects of iron metabolism, including iron transport, storage, and heme synthesis^4,5^, its most widely accepted and clearly defined role is in the iron-sulfur (Fe-S) cluster biogenesis. Fe-S clusters are inorganic protein co-factors composed of iron and sulfur atoms, with the most common forms being Fe_2_S_2_, Fe_4_S_4,_ and Fe_3_S_4,_ and are essential for the activity of a wide array of proteins involved in key cellular processes including central metabolism, mitochondrial respiration, ribosome biogenesis, DNA replication and repair^6,7^.

In eukaryotes, *de novo* Fe-S cluster biogenesis occurs in mitochondria and relies on a conserved core machinery. This machinery, the ISC complex, comprises NFS1 (a cysteine desulfurase), ISD11, ACP, ISCU (a scaffold protein), and FXN^8–11^ (Figure 1A). The process begins with sulfur mobilization from L-cysteine by NFS1, which generate a persulfide intermediate that is transferred to a conserved cysteine on ISCU. ISD11 is specific to the eukaryotic system and stabilizes NFS1. The iron source is still debated. A mitochondrial ferredoxin/ferrodoxin reductase pair (FDX2-FDXR) provides electrons to reduce the persulfide to sulfide, forming a Fe_1_S_1_ intermediate on ISCU^12^. FXN interacts directly with the ISC complex and acts as a crucial activator, accelerating persulfide formation and transfer^9,13–15^. Subsequent dimerization of ISCU will allow Fe_2_S_2_ formation. Following assembly, Fe-S clusters are stabilized and transferred to target apo-proteins via a transfer machinery including the chaperone (HSPA9), co-chaperone (HSC20), and glutaredoxin GLRX5^12^. In eukaryotes, a degree of complexity is added since Fe-S clusters need to be distributed both to mitochondrial apo-proteins and exported to the cytosol and nucleus^9–11^. For mitochondrial targets, Fe_2_S_2_ clusters may be directly incorporated or further processed via the ISCA complex to generate Fe_4_S_4_ clusters^9–11^ (Figure 1A). Additionally, a iron-sulfur-based compound, herein referred to as (Fe-S)intermediate, (Fe-S)int, synthetized by the ISC complex, is thought to be exported by the mitochondrial ABC transporter ABCB7 to support the Cytosolic Iron Sulfur Assembly (CIA) machinery, enabling the maturation of cytosolic and nuclear Fe-S proteins^9,10,16^ (Figure 1A). Given the nature of Fe-S clusters, iron metabolism plays a central role in regulating their production. However, the mechanisms governing the regulation of Fe-S cluster biogenesis and the distribution of these cofactors among cellular compartments and recipient proteins remain poorly understood.

**Figure 1:**
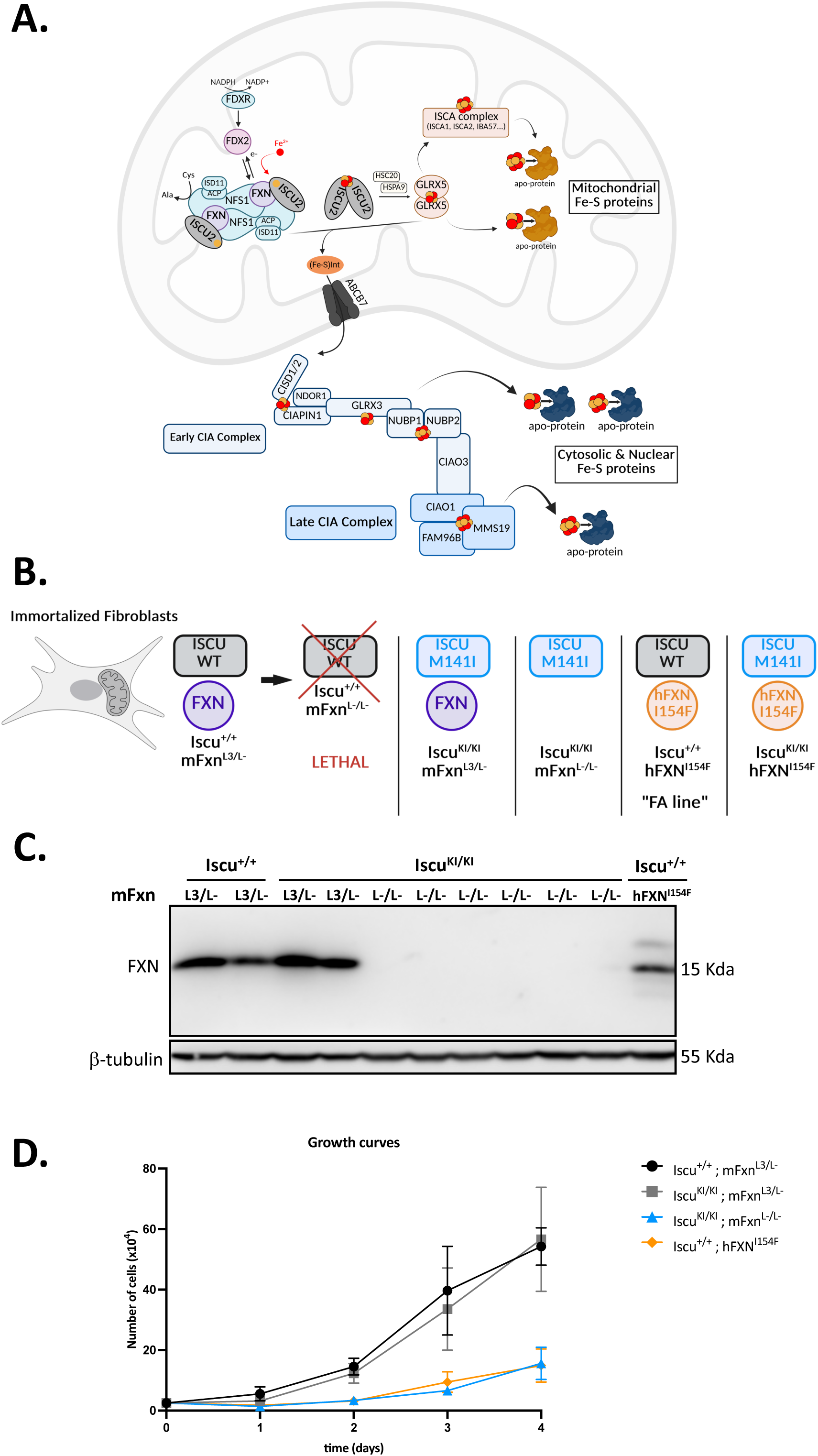
ISCU M141I substitution bypasses lethality induced by absence of frataxin in dividing mammalian cells. (A) Schematic representation of Fe-S cluster biogenesis. In mitochondria, the ISC complex synthetize Fe-S clusters *de novo*. NFS1-ACP-ISD11= sub-complex composed of a cysteine desulfurase providing sulfur to the scaffold for Fe-S cluster synthesis. ISCU= scaffold protein. FXN = regulator, activator of the complex. FDX2-FDXR= ferredoxin/ferrodoxin reductase pair providing electrons for persulfide to sulfide reduction. Dimerization of ISCU allows Fe_2_S_2_ formation. HSPA9 = chaperone, HSC20 = co-chaperone, GLRX5 = glutaredoxins are involved in stabilization and transfer of the Fe-S cluster to target apo-proteins and to the cytosolic CIA machinery. ISCA complex= involved in transfer and formation of Fe_4_S_4_ clusters for mitochondrial apo-proteins. Sulfur/Iron based intermediate (FeS)_int._ synthetized by the ISC complex is exported by the mitochondrial ABC transporter ABCB7 to the Cytosolic Iron sulfur Assembly (CIA) machinery. Early CIA complex: CISD1/2, CIAPIN1, GLRX3, NUBP1, NUBP2, CIAO3. Involved in transfer and insertion of Fe_2_S_2_ and Fe_4_S_4_ clusters into a subset of cytosolic and nuclear apo-proteins. Late CIA complex: CIAO1, FAM96B, MMS19. Involved in transfer of Fe_4_S_4_ clusters to a subset of cytosolic and nuclear apo-proteins (mainly involved in genome integrity). (B) Schematic representation of the genotype of all cell lines used in this study. All cells were derived from immortalized murine fibroblasts Iscu^+/+^; mFxn^L3/L-^. (C) Western blot analysis of FXN in Iscu^+/+^; mFxn^L3/L-^ control cells, Iscu^KI/KI^; mFxn^L3/L-^ cells, Iscu^KI/KI^; mFxn^L-/L-^ cells bypassing lethality and Iscu^+/+^; hFXN^I154F^ cells (FA line). β-tubulin was used as a loading control. (D) Growth curves of Iscu^+/+^; mFxn^L3/L-^ control cells, Iscu^KI/KI^; mFxn^L3/L-^ cells, Iscu^KI/KI^; mFxn^L-/L-^ cells bypassing lethality and Iscu^+/+^; hFXN^I154F^ cells (FA line). Cells were plated on day 0 at a density of 2.5×10^4^ cells per well and counted every day for four days. Each point represents the mean of three independent biological experiments.

One of the major differences between eukaryotes and prokaryotes lies in the dependency of Fe-S clusters biogenesis on FXN. In eukaryotes, FXN plays an essential role, as its complete absence is embryonic lethal in mouse and causes severe growth defects in the yeast *Saccharomyces cerevisiae*^17,18^. In the context of FA, reduced FXN levels impair Fe-S cluster biogenesis, leading to a deficit in Fe-S dependent proteins. This deficit contributes to the main hallmarks of FA, which includes pronounced mitochondrial dysfunction, dysregulated iron homeostasis characterized by mitochondrial iron accumulation, and increased sensitivity to oxidative stress^19–21^. In contrast, in some bacteria such as *E. coli*, Fe-S cluster biogenesis is independent of the frataxin homolog CyaY. Indeed, complete deletion of *cyaY* in *E. coli* results in no observable phenotypes^22,23^. The molecular basis underlying this differential requirement of FXN remains unclear, however, studies in *E. coli* and *S. cerevisiae* have identified a single amino acid substitution in the scaffold protein that can alter FXN dependency. In yeast, a methionine to isoleucine at position 141(M141I) in the main scaffold protein Isu1 was identified in spontaneous suppressor strains that restored growth in frataxin-deficient background^24^. These strains also showed rescue of iron homeostasis and mitochondrial Fe-S protein activities^25,26^. Intriguingly, in *E. coli*, the equivalent position (residue 108) of IscU naturally contains isoleucine. Replacing this isoleucine with methionine (I108M) rendered *E. coli* dependent on CyaY, with growth defects and deficits in Fe-S proteins, mimicking eukaryotic FXN dependence^27^. Biochemical studies suggest that the M141I mutation may enhance cysteine desulfurase activity^25^, while other *in vitro* work has shown that the human ISCU M140I variant increases the rate of Fe-S cluster transfer to GLRX5, implying a possible role in the downstream steps of the pathway^28^. However, whether such bypass mechanisms are effective in mammalian cells or *in vivo* has not been explored.

To address this, we introduced the ISCU M141I variant in murine fibroblasts carrying the conditional *Fxn* allele^29^ using CRISPR-Cas9 mediated genome editing on the endogenous Iscu gene. We found that the resulting ISCU M141I substitution enabled cell survival in the absence of FXN, supporting proliferation. However, surviving clones exhibited reduced growth rates, persisting mitochondrial dysfunction, and deficits in mitochondrial Fe-S protein activities. Remarkably, cytosolic and nuclear Fe-S proteins remained functional, with normal cell cycle progression and reduced basal DNA damage. Furthermore, we generated a knock-in mouse model harboring the ISCU M141I variant. In this context, the M141I substitution was sufficient to delay the embryonic lethality due to FXN loss, extending viability to at least embryonic day 8.5 (E8.5). These findings provide the first *in vivo* evidence that a single amino acid substitution in the ISCU scaffold protein can partially bypass the requirement for FXN in mammals. They also raise important questions about the rate of Fe-S cluster biogenesis in ISCU M141I cells and the compartmental regulation of Fe-S cluster biogenesis.

## Materials and Methods

### Cell Culture Condition

All the cell lines used in this study were built from the NC6 (Iscu +/+; mFxn L3/L-) immortalized murine fibroblast lines, previously established from frataxin heterozygous mice (mFxn L3/L-)^29^. (Iscu KI/KI; mFxn L3/L-) cell lines expressing ISCU M141I variant were obtained by introduction of a single substitution in the endogenous Iscu gene (see procedure below). (Iscu KI/KI; mFxn L-/L-) cell lines were obtained by expression of a Cre-recombinase (see procedure below). (Iscu +/+; hI154F) were generated in the previous study^29^. Fibroblast lines were grown in DMEM media 1 g/L glucose (Gibco^TM^ 31885084) supplemented with 10% Fetal Bovine Serum (Gibco^TM^ 10500064) and either 50 μg/ml Gentamycin or 100 μg/ml penicillin/streptomycin. Cells were passaged every two or three days using Trypsin/EDTA 0,05% (Gibco^TM^ 25300096).

### Engineering of (Iscu KI/KI; mFxn L3/L-) cells by CRISPR-Cas9

A CRISPR-Cas9 system combined to a homology-directed repair template was designed to introduce a G to C substitution in the Iscu gene to convert the methionine 141 into an isoleucine. The pX330-Cas9-T2A-GFP vector was used to express *Streptococcus pyogenes* Cas9 along with a GFP protein and the guide RNA. Sequence of the sgIscu-1 guide RNA (5’-CGTCTTCTGCCAGCACTGGA-3’) was cloned in the pX330-Cas9-T2A-GFP vector under a U6 promoter using the BsbI restriction site. The sgIscu-1 guide RNA was designed to target a region in exon 5 of the Iscu gene near the ATG coding for the methionine 141. The repair template was designed as a long single-strand oligonucleotide ssODN (5’CAATGCTGTATGTGTGTAAGTGTATAAGATGTAATAAAGACCATGCCTCTGGTGTCAGAC ATTGAAGCTGTCTACCTTCCCATGCCTTCCAGT**C**CT**C**GC**T**GA**G**GA**T**GCCATCAAGGCCGCCC TGGCTGACTACAAACTGAAGCAAGAGTCCAAGAAGGAGGAGCCAGAGAAGCAGTGAGCCCTGGAGACACTCCAGCCA 3’) homologous to the exon 5 region of the Iscu gene with the exception of the G to C substitution and four other silent substitutions which create a EcoNI restriction site for screening (Supp Figure 1B). NC6 L3/L-immortalized murine fibroblast lines were co-transfected with 2-10 μg of pX330-sgIscu-1-Cas9-T2A-GFP vector and 500, 1000, 1500 or 3000 ng of ssODN repair template using lipofectamine^TM^ 2000 (Thermo Fisher 11668019) according to manufacturer’s protocol. Cell sorting of GFP positive cells was performed 48h post-transfection using a BD FACSAria^TM^ II Cell Sorter (Biosciences) and put back in culture at one cell per well in 96-well culture plates. After clonal amplification, PCR-screening using Screen2_Iscu_Fd (5’-GCACATTCAGCCACTCACAT-3’) and Screen2_Iscu_Rv (5’-TAGTGTGCGTTTTGCGTAGC-3’) primers and followed by EcoNI digestion was realized to identify recombined clones carrying the substitution ISCU M141I (Iscu^KI/KI^; mFxn^L3/L-^). Genotype of the obtained clones was additionally confirmed by Sanger sequencing. A total of four (Iscu^KI/KI^; mFxn^L3/L-^) clones were obtained.

### Deletion of FXN in (Iscu KI/KI; mFxn L3/L-) cells by Cre-recombinase

Two (Iscu KI/KI; mFxn L3/L-) clones were transfected with a pEGFP-C2-Cre vector, expressing the bacteriophage P1 Cre recombinase^29^, using lipofectamine^TM^ 2000 (Thermo Fisher 11668019) according to manufacturer’s protocol. Cell sorting of GFP positive cells was performed 48h post-transfection using a BD FACSAria^TM^ II Cell Sorter (Biosciences) and put back in culture at one cell per well in 96-well culture plates. After clonal amplification, FXN deletion was analysed by PCR to identify the (Iscu^KI/KI^; mFxn^L-/L-)^ clones. A total of eight (Iscu KI/KI; mFxn L3/L-) clones were obtained. Experiments conducted in this study were performed with two to three of them.

### Protein extracts and Western Blots

Fibroblasts were collected by Trypsin/EDTA and centrifuged at 1000 rpm for 5 min. Pellets were washed once with Phosphate Buffer Saline (PBS) 1X. Pellets were resuspended in RIPA lysis buffer (Tris HCl pH8 50mM, NaCl 150 mM, sodium deoxycholate 0.5%, SDS 0.1%, Triton X-100 1%, NP-40 1%) supplemented with phosphatase/protease inhibitors and incubated 30 min on ice, then centrifuge at 13 000 rpm at 4°C for 20 min. Supernatant was collected and protein concentration measured by Bradford Assay. Total protein extracts (10 to 20 μg) were pre-stained for 30 min using Cy5-QuickStain Protein Labelling Kit (Amersham^TM^ RPN4000) according to manufacturer’s instruction before being mixed with loading buffer and boiled for 5 min. Proteins were then loaded and separated onto SDS tris-glycine polyacrylamide gels followed by a tank-transfer onto a nitrocellulose membranes. After transfer, total protein Cy5 fluorescence was acquired on a ChemiDoc^TM^ Touch Imaging System (Biorad). Membranes were then blocked for 1h in Tween-20 (0,05%) Tris-buffered saline (TBST) containing 5% of non-fat milk. Membranes were incubated overnight at 4°C with the primary antibody (see below). Membranes were then washed 3 times 10 min in TBST and incubated with a secondary antibody coupled to peroxidase (goat anti-rabbit (Cell Signaling 7074) or goat anti-mouse (Cell Signaling 7076), IgG diluted at 1/5 000 or 1/10 000) at room temperature for 1h. Membranes were then washed 3 times 10 min in TBST and signal was revealed by SuperSignal^TM^ West Femto Maximum Sensitivity Substrate (Thermo Scientific^TM^ 34096) using a ChemiDoc^TM^ Touch Imaging System (Biorad).

Antibodies used: Mouse Anti-FXN (Homemade 4F9), Mouse anti-SDHB (Abcam ab14714), Rabbit anti-LA (Calbiochem 437695), Mouse anti-NDUFS3 (Invitrogen 439200 17D95), Mouse anti-PDCE2 (Santa Cruz sc271352 C-9), Rabbit anti-LIAS (Abcam ab96302), Mouse anti-NFS1 (Santa Cruz sc365308 B7), Rabbit anti-FDX (Homemade 3037), Rabbit anti-ABCB7 (Homemade PJ173), Mouse anti-βtubulin (Invitrogen MA516308)(IGBMC 2A2), Mouse anti-GAPDH (Milipore MAB374 6C5), Rabbit anti-FECH (Proteintech 14466-1-AP), Rabbit anti-NFU1 and anti-ISCU (Covance custom-made), Rabbit anti-POLD1 (Proteintech 15646-1-AP), Rabbit anti-RTEL1 (Proteintech 25337-1-AP), Rabbit anti-ELP3 (Abcam ab190907), Rabbit anti-DPYD (Proteintech 27662-1-AP), Rabbit anti-PPAT (Proteintech 15401-1-AP), Rabbit anti-GLRX3 (Proteintech 11254-1-AP), Mouse anti-β-Actin (Sigma A2228), Mouse anti-FLAG (Origene Technologies TA50011), Mouse anti-SDHA (Abcam ab14715), Rabbit anti-MMS19 (Proteintech 16015-1-AP), Mouse anti-CIAO1 (Santa-Cruz Biotechnology 374498), Rabbit anti-FAM96B (Proteintech 20108-1-AP).

### Enzymatic activities measurements

#### SDH activity measurment

Succinate dehydrogenase (SDH) and isocitrate dehydrogenase (IDH) activities were measured as previously described^30^ using a Cary50 spectrophotometer (Varian SA). Specific activity of SDH (which contains three Fe-S clusters) was normalized against the specific activity of IDH (which does not contain Fe-S clusters).

#### DPYD activity assay

The dihydropyrimidine dehydrogenase (DPYD) activity was determined by thin layer chromatography (TLC), following a previously described protocol (PMID: 34083449; 38950322). Briefly, cell lysates containing 150 μg of proteins isolated from control or Iscu^KI/KI^ cell lines, as specified in the main text and figure legend, were applied to 50 μl of a reaction mix containing 25 mM Tris-HCl (pH 7.5), 0.1% digitonin, 2.5 mM MgCl_2_, 2mM DTT, 10 μM [4-^14^C]-thymine (0.1mCi/ml Moravek Inc. CA, USA), 10 mM NADPH. After 4 hours of incubation at 32 °C, the reaction was stopped by addition of 10 μl of perchloric acid (10% v/v). Reaction mixtures were centrifuged at 20000 x g for 5 minutes and the supernatants analyzed by TLC. Iron Responsive Element (IRE) binding activities of IRP1 and IRP2

The IRE-binding activities of IRPs were analyzed by biotinylated ferritin H (FTH) IRE pulldown, as previously described (PMID: 38732071). The FTH IRE probe was labeled by 3’-biotinylation and ordered from Integrated DNA Technologies.

The sequence is as follows: 5′-GGAGUUCCUGCUUCAACAGUGCUUGGACGGAACUCC-3′. Cell lysates were prepared in lysis buffer (40 mM KCl, 25 mM Tris-Cl, pH 7.5, 1% Triton X-100) supplemented with 1x Halt^TM^ Protease Inhibitor Cocktail (ThermoFisher Scientific, Catalog No: 78430), 1 mM DTT, and 20u/ml RNase Inhibitor (ThermoFisher Scientific, Catalog No: AM2696). Five pmols of annealed IRE probe was incubated with 100 μg of DynaBeads M280 for 20 min at room temperature on a shaker to allow formation of the probe/bead complexes. Following three washes with lysis buffer, the probe/bead complexes were added to 100 μg lysates and incubated at room temperature on a shaker for 20 min. The probe/bead/IRPs complexes were isolated with a magnet and washed with lysis buffer three times prior to being resuspended in 100 μL of 1x NuPage LDS loading buffer. The samples were denatured at 70 C for 5 min. 25 μL of eluates were loaded onto 4-12% Bis-Tris gels for immunoblots against IRP1 and IRP2 (both antibodies are custom made at Covance).

### Growth curve

Cells were plated in a 12-well cell culture plate at 2.5×10^4^ cells per well on day 0. Cells were collected from two wells using Tryspin/EDTA 0,05% (Gibco^TM^ 25300096) each day for four days and counted independently using a Malassez chamber. The mean between the two wells were calculated to have the number of cells at each time points. This experiment was repeated three times independently. Growth curves were realized by ploting the mean of the three independent experiments for each time points.

### Cell cycle analysis

Cells in exponential phase were harvested and fixed with cold 70% ethanol for one hour at 4°C, then centrifuge for 2 min at 4 000 rpm. Pellets were resuspend and incubate in a solution of Phosphate Buffer Saline (PBS)- Triton X-100 0,25% for 15 min at 4°C. Cells were again centrifuge for 2 min at 4 000 rpm, then pellets were resuspend and incubate in a solution of PBS-RNAse A 10 μg/ml-Propidium iodide 20 μg/ml for 30 min at room temperature in the dark. Propidium iodide fluorescence was analyze using a BD FACS CantoII (Bioscience).

### Gamma γH2AX immunofluorescence

Cells were cultivated on coverslips then fixed with 4% paraformaldehyde (PFA) for 15 min at room temperature. After 3 washes with PBS for 5 min, permeabilization was performed in a PBS-Triton 0,1% solution for 10 min at room temperature. Coverslips were washed twice in PBS for 5 min, then blocking was performed in a PBS-BSA 3%-Tween 0,05% solution for 30 min. Coverslips were then incubated with a primary antibody anti-γH2AX (Ser139) (Sigma 05-636, clone JBW301) in half diluted blocking solution for 1h at room temperature. After 3 washes of 10 min in PBS, coverslips were incubated with a goat anti-mouse Alexa Fluor 594 Secondary Antibody (Invitrogen^TM^ A-11032) (dilution 1/1000) and DAPI (dilution 1/1000) for 1h at room temperature in the dark. Coverslips were washed 3 times for 10 min in PBS then mount in AquaPolyMount (Polysciences) and dry overnight. Image acquisition was performed on a Z1 epifluorescence microscope (Zeiss).

### Seahorse Analysis

Oxygen Consumption Rate (OCR) was measured using a Seahorse Analyzer XFe96 (Agilent) and Seahorse XF Cell Mito Stress Test Kit (Agilent 103015-100). Between 12 000 and 6 000 cells per well, depending on the cell line, were seeded in 96-well Seahorse plate (Agilent 103794-100). The next day, Seahorse XF Cell Mito Stress Test Kit protocol was followed according to manufacturer’s instructions using the Seahorse XF DMEM media supplemented with 1 mM pyruvate, 2 mM glutamine, 10 mM glucose (Agilent 103680-100) and the following optimized concentrations of respiratory chain inhibitors: 1.5 µM Oligomycin, 1.5 µM FCCP and 0.5 µM Rotenone/Antimycin A. Mixing steps (Wave Controller 2.4) were reduced to 60 seconds to prevent detaching of the cells from the bottom of the well. After Seahorse measurements, 150 µl of media per well were removed, then 100 µl per well of fixation solution (PFA 4%, Triton 0.2% and DAPI 1/500) were add. Cells were fixed for 15 min at room temperature in the dark, then washed 3 times 5 min with PBS. Then, a picture of each well was performed on an fluorescence microscope (EVOS M5000) and the number of cells per well were count based on DAPI staining using a homemade macro on Fiji/ImageJ software. OCR values measured by Seahorse were normalized by the number of cells per well.

### Radioactive iron incorporation assay

The ^55^Fe incorporation assays into C-terminally FLAG-tagged POLD1 or DPYD transiently expressed in control and Iscu^KI/KI^ cell lines were performed as previously described (PMID ID: 34083449; 29309586). Briefly, cells were grown in expression medium in the presence of 1 μM ^55^Fe-Transferrin. Cytosolic extracts were subjected to immunoprecipitation with anti-FLAG to immunocapture FLAG-tagged proteins 48h post-transfection. Samples collected after competitive elution (with 3X FLAG peptide at 100μg/ml) were analyzed by scintillation counting to assess ^55^Fe content. The background levels corresponding to ^55^Fe measurements on eluates after anti-FLAG immunoprecipitations on cytosolic extracts isolated from cells transfected with the empty vector, pCMV6-Entry (Origene Technologies), were also included to account for nonspecific ^55^Fe amounts stochastically associated to the beads. In parallel, 10 µL of eluates were run on 4-16% native PAGE gels (ThermoFisher), resolved after run at 150V for 1 hour, followed by 2 hours at 250V, and imaged by autoradiogram.

### Mitochondrial Iron Measurment

Subcellular fractionation into cytosol and intact mitochondria was done as previously described (PMID: 24606901; 26749241). Briefly, mitochondria from cell pellets (∼10^9^ cells) were isolated from the cytosolic fractions after cell permeabilization with a buffer containing 0.1% digitonin in 210 mM mannitol, 20 mM sucrose, and 4 mM HEPES. The pellets after centrifugation at 700 x *g* for 5 min contained mitochondria, which were isolated by differential centrifugation and solubilized in lysis buffer I, containing 50 mM Bis-Tris, 50 mM NaCl, 10% w/v Glycerol, 0.001% Ponceau S, 1% lauryl maltoside, pH 7.2 and protease inhibitors.

Total iron content on isolated mitochondria was measured by Inductively Coupled Plasma Mass Spectrometry (ICP-MS) (Agilent model 7900). For each sample, 200 µL of concentrated trace-metal-grade nitric acid (Fisher Scientific) was added to 200 µL of sample taken in a 15 mL Falcon tube. The tubes were then incubated at 85 C in an oven for overnight digestion. Each sample was diluted to a total volume of 4 mL with deionized water, and analyzed by ICP-MS. Iron content was normalized by protein concentrations on mitochondrial extracts.

### Iscu^KI/KI^ and Iscu^KI/KI^; mFxn^+/L^ Mice generation and breeding

The M141I point mutation in the Iscu gene was introduce into the mouse genome of C57BL/6N fertilized eggs using a CRISPR-Cas9 system combined to a homology-directed repair template (HDR) A 124 nts single strand DNA (GTAATAAAGACCATGCCTCTGGTGTCAGACATTGAAGCTGTCTACCTTCCCATGCCTTCCAG T<u>C</u>CT<u>C</u>GC<u>G</u>GAAGACGCCATCAAGGCCGCCCTGGCTGACTACAAACTGAAGCAAGAGTCCAAG was coelectroporated with 1.2µM of spCas9 ((Alt-R SpCas9 Nuclease V3; ref 1081059)) and 6 µM of crRNA (CGTCTTCTGCCAGCACTGGA) and tracRNA (IDT). To facilitate genotyping and avoid new double strand break once the HDR event occurred, a BstUI restriction site was engineered into the repair template alongside the mutation. The presence of the expected G>C mutation was confirmed by Sanger sequence on aach BstUI positive F1 heterozygous animal. DNA from tail biopsies was amplified by PCR around the point mutation insertion site on the Iscu gene. PCR products were digested with BstUI to distinguish between wild-type, heterozygous and homozygous alleles based on the presence or absence of the restriction site. Digestion products were resolved by agarose gel electrophoresis to determine genotype. Selected founders were bred in order to obtain germ line transmission of the mutation.

The compound line *Iscu^KI/KI^; mFxn^+/L^* was generated by crossing *Iscu^KI/KI^* homozygous mice with heterozygous (*mFxn^+/L-^*) mice from the DA16 line^30^. Genotypes of the offsprings were determined from DNA of tail biopsies. The same procedure as describe before was used to verify the presence of the point mutation on the Iscu gene. Genotyping of the mFxn gene was performed by two PCR amplifications using either a set of primers which amplifies a region in exon 4 corresponding to a wild-type allele (+) or a set of primers that amplifies a region corresponding to an allele with a deleted exon 4 (L-).

Mice were maintained in a pathogen-free animal facility with a 12-hour light/dark cycle and provided with standard rodent chow (D04, SAFE, Villemoisson-sur-Orge, France), water and libitum. All animal procedures were performed in accordance with institutional animal care guidelines and approved by the local ethical committee (APAFIS# 54248-2024102512328818).

## Results

### ISCU M141I substitution bypasses lethality induced by absence of frataxin in dividing mammalian cells

We utilized previously described immortalized mouse fibroblasts^29^, which carry an allele deleted for exon 4 (50% expression of frataxin) and a conditional Fxn allele with LoxP-flanked exon 4. Cre-mediated excision of the conditional allele leads to complete loss of FXN protein expression (Figure 1B)(Supp Figure 1A) and consequent cell cycle arrest within 3 days, ultimately resulting in cell death^29^ (Figure 1B). To investigate whether the ISCU M141I variant may exert an effect on this phenotype, the substitution was introduced on the endogenous Iscu locus using a CRISPR-Cas9 genome editing combined with a homology-directed repair template (Supp Figure 1B). Following clonal selection and validation by PCR followed by enzymatic digestion (Supp Figure 1C) and sequencing, four independent homozygous *Iscu^KI/KI^; mFxn^L3/L-^* clones were obtained (Figure 1B). These clones displayed similar FXN protein levels and proliferation rates compared to control (*Iscu^+/+^, mFxn^L3/L-^*) cells (Figure 1C and D). Introduction of the M141I mutation led to a slight reduction in ISCU protein levels without affecting the expression of other ISC core components such as NFS1 or FDX (Supp Figure 2A).

**Figure 2:**
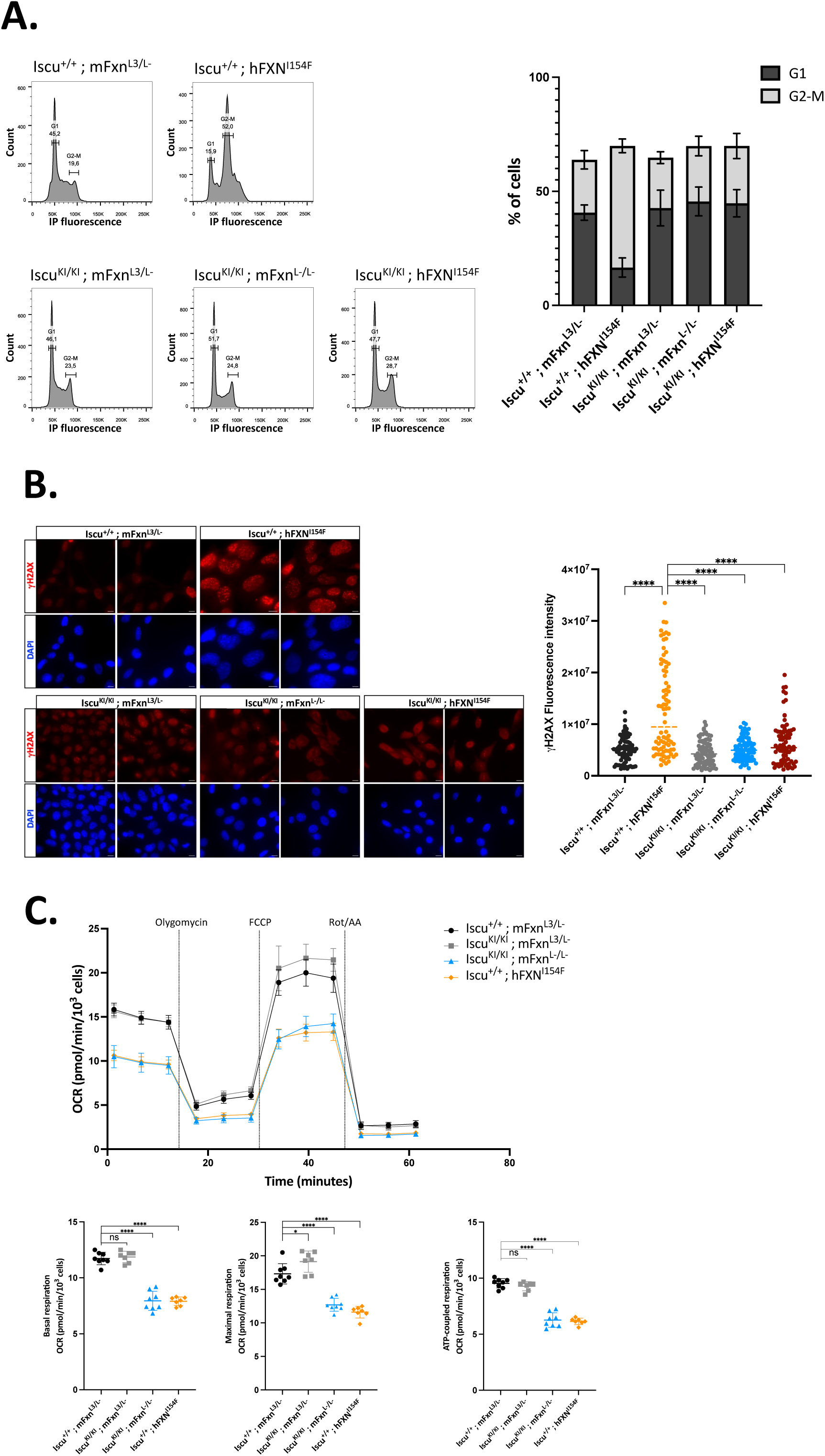
ISCU M141I prevents cell cycle arrest and DNA damage accumulation, but mitochondrial dysfunction persists in the absence of FXN protein. (A) Cell cycle progression analysis assessed by propidium iodide staining and flow cytometry. Representative flow cytometry graphics from three independent biological experiments are shown. Bar graphs display the percentage of cells in G1 and G2/M phases, presented as the mean ± standard deviation of three independent experiments. (B) Representative images of γH2AX immunofluorescence (red) and DAPI staining (blue). The accompanying graph represents the quantification of γH2AX immunofluorescence intensity, each point represents a single nucleus. A total of 80 nuclei for each cell line were analyzed. Statistical significance was assessed by one-way ANOVA (**** p<0.0001). (C) Oxygen Consumption Rate (OCR) was measured using Seahorse Analyzer following sequential addition of the mitochondrial inhibitors oligomycin, FCCP and rotenone/antimycin A (Rot/AA). For each cell lines, OCR was measured in 8 technical replicates. Data are normalized to the cell number and expressed as pmol/min/10^3^ cells. Lower panels show quantification of basal respiration, maximal respiration and ATP-coupled respiration, derived from OCR traces. Statistical comparisons were performed using one-way ANOVA (**** p<0.0001)(* p<0.05). Shown is a representative experiment out of three biological replicates.

FXN deletion was subsequently introduced in two independent *Iscu^KI/KI^; mFxn^L3/L-^* clones via transient transfection of a pCre-EGFP plasmid (Supp Figure 1A). Clonal selection yielded viable *Iscu^KI/KI^; mFxn^L-/L-^* cells (Figure 1B), which lacked detectable FXN protein, as assessed by western blot (Figure 1C), demonstrating that ISCU M141I variant allows to bypass lethality due to FXN absence in dividing cells (Figure 1B). FXN deletion in these cells did not change protein levels of the components of the ISC machinery core, including ISCU (Supp Figure 2A). Despite bypassing lethality, the viable *Iscu^KI/KI^; mFxn^L-/L^* cells exhibited significantly slower proliferation rate relative to both parental *Iscu^KI/KI^; mFxn^L3/L-^* and control *Iscu^+/+^; mFxn^L3/L-^*lines (Figure 1D), suggesting that ISCU M141I permits survival.

### Establishing an appropriate control to investigate the effect of ISCU M141I variant on Fe-S cluster biogenesis

Our first challenge in investigating the effect of the ISCU M141I variant on Fe-S cluster biogenesis, in the presence or absence of FXN, was selecting an appropriate control. As FXN-null (*Iscu^+/+^; mFxn^L-/L-^*) dividing cells are not viable in culture, we employed an established FA patient-like cell line expressing the pathogenic human FXN I154F variant (*Iscu^+/+^; hFXN^I154F^*) with a complete deletion of mouse frataxin as a functional comparator^29^ (Figure 1B)(Supp Figure 1A). The I154F point mutation, found in FA patients, impairs FXN processing, resulting in low levels of mature FXN protein^31^. Western blot analysis confirmed the accumulation of the FXN intermediate form at 19 kDa in the *Iscu^+/+^; hFXN^I154F^* cell line (Figure 1C). Biochemical evidence also suggests that this point mutation might decrease the activity of the cysteine desulfurase NFS1^32^. The *Iscu^+/+^; hFXN^I154F^* cell line has been shown to exhibit impaired Fe-S cluster biogenesis, resulting in mitochondrial dysfunction and other characteristic hallmarks of FA, such as iron dysregulation and increased sensitivity to oxidative stress^29^. Notably, this cell line displays slow cellular growth, comparable to that observed in the viable *Iscu^KI/KI^; mFxn^L-/L-^* cells (Figure 1D). We will refer to the *Iscu^+/+^; hFXN^I154F^*line as the “FA line” throughout the manuscript.

### ISCU M141I prevents cell cycle arrest and DNA damage accumulation, but mitochondrial dysfunction persists in absence of FXN protein

To investigate the basis of the reduced growth observed in *Iscu^KI/KI^; mFxn^L-/L-^* cells bypassing lethality due to FXN loss, we first assessed cell proliferation and metabolic state. Cell cycle progression was analyzed using propidium iodide staining and flow cytometry to quantify nuclear DNA content in all cell lines. Both control *Iscu^+/+^; mFxn^L3/L-^* and *Iscu^KI/KI^; mFxn^L3/L-^* lines displayed a typical profile of proliferating cells, with 40.7% (± 3.3%) of cells in G1 and 23.2% (± 4.1%) in G2/M (Figure 2A). In contrast, the FA line (*Iscu^+/+^; hFXN^I154F^*) showed a marked G2/M arrest (∼53.3% ±3.0%), indicative of impaired cell division (Figure 2A). This G2/M block likely results from defective Fe-S cluster biogenesis affecting nuclear Fe-S dependent proteins involved in DNA replication and repair. Notably, expression of the ISCU M141I variant in the absence of FXN protein (*Iscu^KI/KI^; mFxn^L-/L-^*), prevented G2/M arrest, with cell cycle distribution resembling control cells (Figure 2A). This indicates that ISCU M141I variant prevents cell cycle blockade due to FXN loss. Interestingly, when the mutant human FXN I154F protein was expressed in *Iscu^KI/KI^; mFxn^L-/L-^*cells to generate the *Iscu^KI/KI^; hFXN^I154F^*line (Figure 1B)(Supp Figure 1A), no G2/M cell cycle arrest was observed suggesting that ISCU M141I is epistatic on FXN I154F (Figure 2A).

Given the link between G2/M and DNA repair deficiency, we next assessed DNA damage by γH2AX immunofluorescence. FA line (*Iscu^+/+^; hFXN^I154F^*) showed elevated γH2AX foci relative to control cells (*Iscu^+/+^; mFxn^L3/L-^* or *Iscu^KI/KI^; mFxn^L3/L-^*) (Figure 2B). In contrast, *Iscu^KI/KI^; mFxn^L-/L-^* exhibited γH2AX levels comparable to controls (Figure 2B), indicating reduced DNA damage accumulation. Similarly, *Iscu^KI/KI^; hFXN^I154F^* cells showed no increase in γH2AX (Figure 2B), further confirming that the ISCU M141I variant preserves genome integrity in the absence of FXN. These findings suggest that ISCU M141I restores Fe-S dependent DNA metabolism pathways, preventing cell cycle arrest and genome instability despite FXN deficiency. However, the continued slow growth of *Iscu^KI/KI^; mFxn^L-/L-^* cells prompted further investigation into mitochondrial function.

We evaluated mitochondrial respiration using Seahorse assays, measuring oxygen consumption rates (OCRs) following sequential addition of oligomycin, FCCP and rotenone/antimycin A. In line with the previously described mitochondrial dysfunction^29^, the FA line showed reduced basal, maximal, and ATP-coupled respiration (Figure 2C). No mitochondrial defects were observed in *Iscu^+/+^; mFxn^L3/L-^* or *Iscu^KI/KI^; mFxn^L3/L-^* cells, indicating that ISCU M141I does not impair mitochondrial respiration in the presence of FXN (Figure 2C). However, *Iscu^KI/KI^; mFxn^L-/L-^* cells exhibited significant reductions in basal, maximal and ATP-coupled respiration, to the same extent as the FA line (Figure 2C). Moreover, ultrastructural analysis revealed mitochondrial defects, including mitochondria with reduced matrix density, cristae disorganization, and increased autophagosome formation (Supp Figure 3). These results demonstrate that ISCU M141I cannot rescue the mitochondrial dysfunction caused by FXN loss.

**Figure 3:**
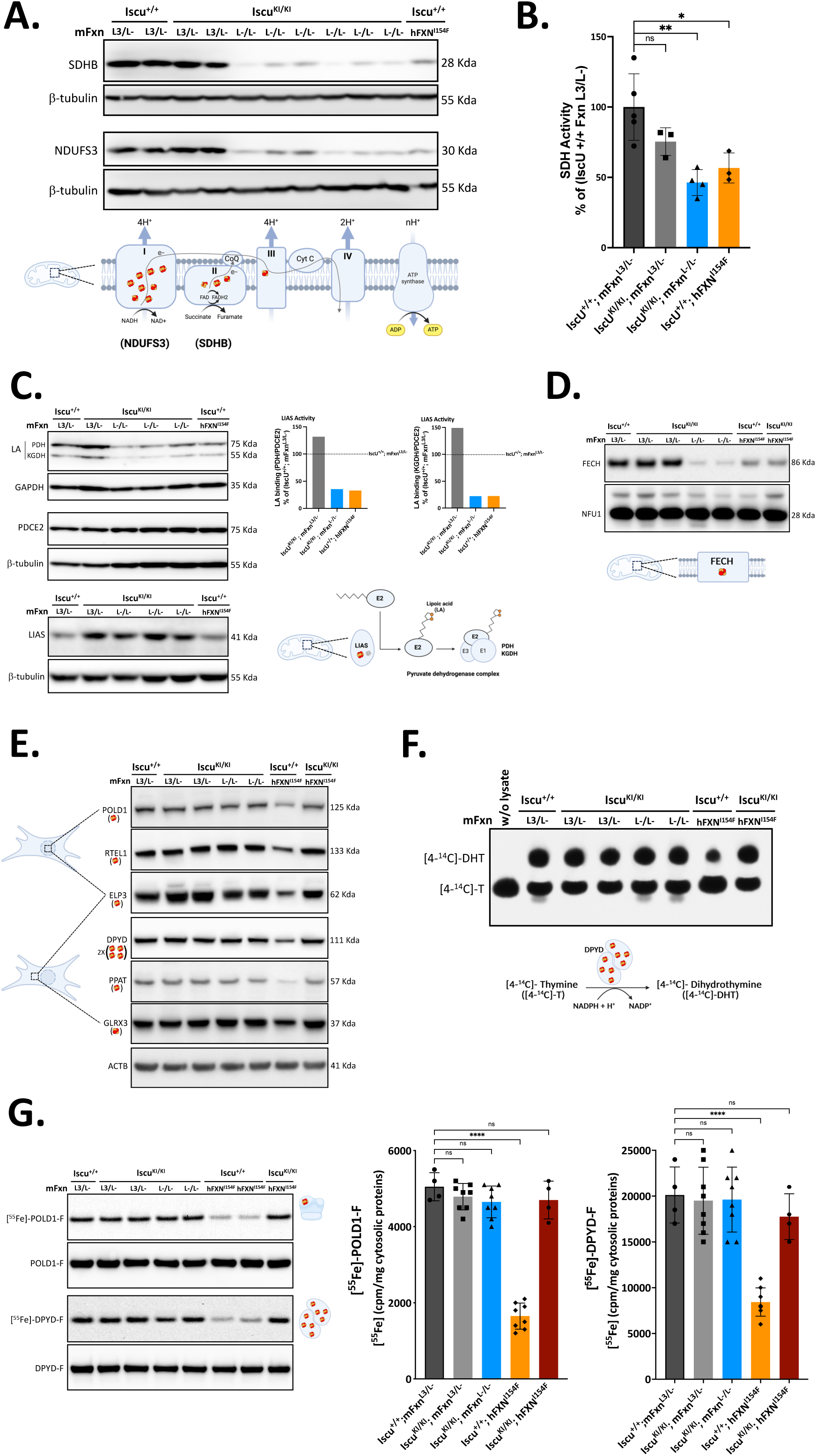
ISCU M141I does not compensate for FXN loss in supporting Fe-S cluster biogenesis for mitochondrial apo-proteins but restores Fe-S biosynthesis for [or incorporation into] nuclear and cytosolic apo-protein. (A) Western blot analysis of mitochondrial Fe-S cluster containing subunits of the respiratory chain. SDHB= subunit of complex II, NDFUS3= subunit of complex I. (B) Spectrophotometric measurements of succinate dehydrogenase (SDH) enzymatic activity, normalized to isocitrate dehydrogenase (IDH). SDH activity is expressed as a percentage relative to Iscu^+/+^; mFxn^L3/L-^ control cells activity. Iscu^+/+^; mFxn^L3/L-^ n=5, Iscu^KI/KI^; mFxn^L3/L-^ n=3, Iscu^KI/KI^; mFxn^L-/L-^ n=4, Iscu^+/+^; hFXN^I154F^ n=5. Statistical comparisons were performed using one-way ANOVA (** p<0.005) (* p<0.05). (C) Representative Western blot (n=3) of lipoic acid (LA) bound to PDH and KGDH as a readout of LIAS activity. Total PDCE2, GAPDH, and β-tubulin are shown as loading controls. LA signal was quantified on PDH and KGDH, normalized to their respective loading control and to total PDCE2. Western blot analysis of total LIAS protein level. (D) Western blot analysis of mitochondrial Fe-S cluster containing protein FECH (Ferrochelatase). (E) Western blot analysis of nuclear and cytosolic Fe-S cluster containing proteins POLD1, RTEL1, ELP3, DPYD, PPAT, and GLRX3. (F) DPYD enzymatic activity assay based on the conversion of [4-^14^C]-thymine ([4– ^14^C]-T) to [4-^14^C]-dihydrothymine ([4–^14^C]-DHT) using thin-layer chromatography and autoradiography. A reaction mix without cell extract was used as a negative control to identify unreacted substrate. (F) ^55^Fe incorporation into C-terminally FLAG-tagged POLD1 (POLD1-F) and DPYD (DPYD-F). ^55^Fe levels were quantified using scintillation counting (n = 4 biological replicates, with two different clones for Iscu^KI/KI^; mFxn^L3/L-^ cells, Iscu^KI/KI^; mFxn^L-/L-^ cells bypassing lethality pooled together, and Iscu^+/+^; hFXN^I154F^ cells (“FA cells”)). Statistical analysis was performed using one-way ANOVA (**** p<0.0001).

Taken together, our data suggest that ISCU M141I variant partially compensates for FXN loss by supporting Fe-S cluster biogenesis sufficient for cytosolic and nuclear Fe-S proteins, thereby maintaining genome stability and cell division, and preventing lethality. However, mitochondrial respiratory defects persist, leading to reduced cell growth rates.

### ISCU M141I does not support Fe-S cluster biogenesis for mitochondrial apo-proteins in the absence of FXN

Building on our previous hypothesis and considering the observed mitochondrial dysfunction, we next sought to assess the ability of the ISCU M141I variant to support Fe-S cluster biogenesis for mitochondrial proteins. Since many Fe-S-containing enzymes require theses co-factors for stability and activity, their loss typically results in decreased protein levels and/or reduced enzymatic function. These properties can be exploited as indirect readouts to evaluate Fe-S cluster biogenesis through western blot and enzymatic assays. Among mitochondrial Fe-S proteins, subunits of respiratory chain complexes I and II are particularly enriched in Fe-S clusters and are known to be prone to degradation when they fail to acquire their cofactors. Complex I contains eight clusters (2 Fe_2_S_2_ and 6 Fe_4_S_4_) distributed across various subunits, while complex II subunit SDHB harbors three clusters (1 Fe_2_S_2_, 1 Fe_3_S_4_, and 1 Fe_4_S_4_). These clusters are delivered by either GLRX5 or the ISCA complex, depending on their complexity (Figure 1A). Consistent with previous findings^29^, we observed decreased levels of both SDHB and the complex I subunit NDUFS3 in FA cells (*Iscu^+/+^; hFXN^I154F^*) (Figure 3A). A similar reduction was found in viable *Iscu^KI/KI^; mFxn^L-/L-^* cells, whereas protein levels remain unchanged in *Iscu^KI/KI^; mFxn^L3/L-^* cells prior to FXN deletion (Figure 3A). To further assess complex II function, we measured succinate dehydrogenase (SDH) activity of complex II, which is directly dependent on the Fe-S cofactors in SDHB. As previously reported^29^, the FA line retained only ∼56.7% (± 10.7%) of SDH activity relative to control lines (Figure 3B). In *Iscu^KI/KI^; mFxn^L-/L-^* cells, SDH activity was similarly reduced to ∼46.3% (± 9.1%) (Figure 3B), in correlation with the decreased SDHB protein levels observed. In contrast, no significant change in SDH activity was observed in *Iscu^KI/KI^; mFxn^L3/L-^* cells prior to FXN loss, although a slight downward trend was noted (Figure 3B). These results indicate that the ISCU M141I variant does not compensate for FXN in supplying sufficient Fe-S clusters to the respiratory chain complexes, which likely underlies the mitochondrial dysfunction observed in FXN-deficient cells.

To determine whether the Fe-S cluster deficiency extended beyond OXPHOS complexes, we analyzed lipoic acid synthetase (LIAS), a mitochondrial enzyme containing two Fe_4_S_4_ clusters, one catalytic and one serving as a sulfur donor. LIAS catalyzes the lipoylation of specific subunits within the pyruvate dehydrogenase (PDH) and α-ketoglutarate dehydrogenase (KGDH) complexes. It is known that loss of the Fe-S cofactors impairs LIAS enzymatic activity. Using an antibody raised against the lipoate modifications of the subunits of the mitochondrial dehydrogenases and normalizing to the E2 subunit of the pyruvate dehydrogenase complex (PDHCE2), we indirectly assessed LIAS activity (Figure 3C). In both the FA line and the viable *Iscu^KI/KI^; mFxn^L-/L-^* cells, lipoic acid incorporation into PDH and KGDH was reduced, indicating impaired LIAS activity (Figure 3C). No significant difference was observed in *Iscu^KI/KI^; mFxn^L3/L-^* cells, indicating preserved LIAS function prior to FXN loss (Figure 3C). Interestingly, we noted an increase in LIAS protein levels in ISCU M141I expressing cells even before FXN deletion, suggesting a compensatory upregulation that becomes insufficient following FXN removal (Figure 3C).

We also assessed ferrochelatase (FECH), an Fe-S protein involved in heme biosynthesis that contains a single Fe_2_S_2_ clusters. FECH protein levels were reduced in the FA line and even more greatly decreased in the *Iscu^KI/KI^; mFxn^L-/L-^* cells (Figure 3D), further supporting the existence of a broad mitochondrial Fe-S cluster defect.

Collectively, these findings demonstrate that the ISCU M141I variant cannot compensate for the absence of FXN in supporting the maturation of mitochondrial Fe-S proteins. The mitochondrial defect observed in the *Iscu^KI/KI^; mFxn^L-/L-^* cells closely mirrors that of the FA line and is independent of the specific Fe-S cluster insertion pathway (ISCA complex vs GLRX5) or the complexity and number of clusters per protein (Figure 1A). This failure to maintain mitochondrial Fe-S proteins likely explains the observed respiratory defects and impaired cell proliferation in the *Iscu^KI/KI^; mFxn^L-/L-^* cells.

### ISCU M141I compensates for FXN deficiency to support Fe-S cluster incorporation into nuclear and cytosolic apo-proteins

To evaluate whether ISCU M141I enables functional Fe-S cluster delivery to cytosolic and nuclear proteins, we investigated the integrity of the Cytosolic Iron sulfur Assembly (CIA) pathway. We began by assessing the expression levels of key CIA components, including the mitochondrial ABC-transporter ABCB7, which is proposed to export a sulfur-containing intermediate (possibly also iron-containing)^33^ generated by the mitochondrial ISC machinery, and the late-acting CIA proteins, MMS19, CIAO1 and FAM96B, which mediate Fe_4_S_4_ cluster transfer to essential nuclear and cytosolic targets (Figure 1A). Western blot analysis revealed no differences in protein levels for ABCB7, MMS19, CIAO1, or FAM96B across all cell lines (Supp Figure 2B), indicating preserved integrity of the proposed export and CIA pathways. Interestingly, GLRX3, a central component of the early CIA machinery that coordinate Fe_2_S_2_ clusters as a homodimer, was selectively decreased in the FA line (*Iscu^+/+^; hFXN^I154F^*) (Figure 3E). Given that GLRX3 stability depends on its ability to dimerize in the presence of a Fe_2_S_2_ cluster^34–36^, its reduction suggests that impaired *de novo* Fe-S cluster synthesis by the ISC machinery, due to the mutant FXN I154F, may limit the availability of the sulfur/iron-based compound for the early CIA machinery. In contrast, GLRX3 protein level remained unchanged in *Iscu^KI/KI^; mFxn^L-/L-^* cells, implying that the ISCU M141I variant supports sufficient export of the sulfur/iron-based compound to sustain early CIA function (Figure 1A).

We then evaluated several late-acting CIA target proteins involved in DNA metabolism, including the catalytic subunit of the DNA polymerase δ (POLD1)(replication/repair, one Fe_4_S_4_), the DNA helicase RTEL1 (telomere maintenance, one Fe_4_S_4_), ELP3 (transcription regulation and tRNA modification, one Fe_4_S_4_), PPAT (purine metabolism, one Fe_4_S_4_) and DPYD (pyrimidine metabolism, eight Fe_4_S_4_). As expected, all five proteins were greatly reduced in the FA line (*Iscu^+/+^; hFXN^I154F^*), in line with the G2/M arrest and compromised Fe-S cluster biogenesis previously observed (Figures 2A and 3E). In contrast, protein levels of these Fe-S proteins were unaffected in cells expressing the ISCU M141I variant, regardless of FXN presence (Figure 3E).

To confirm the functional integrity, we directly measured DPYD activity by monitoring the conversion of [4-^14^C]-thymine to [4-^14^C]-dihydrothymine by thin-layer chromatography (Figure 3F). DPYD activity was significantly decreased in the FA line, consistent with the reduced protein levels, but was preserved in ISCU M141I expressing cells irrespective of FXN status (Figure 3F). We further validated Fe-S incorporation into CIA targets by quantifying radioactive iron [^55^Fe] incorporation into POLD1 and DPYD via scintillation counting and native autoradiography. As expected, a substantial reduction of [^55^Fe] content in both POLD1 and DPYD was observed only in the FA line, whereas ISCU M141I expressing cells, including the *Iscu^KI/KI^; hFXN^I154F^*, retained normal [^55^Fe] incorporation (Figure 3G), reinforcing the conclusion that Fe-S clusters are provided to nuclear and cytosolic apo-protein targets in the presence of the ISCU M141I variant, regardless of FXN presence.

Collectively, these data demonstrate that ISCU M141I variant restores Fe-S cluster delivery to nuclear and cytosolic apo-proteins in the absence of FXN. Interestingly, in the cell line expressing both the ISCU M141I variant and the mutant human FXN I154F (Iscu KI/KI; hI154F), we did not observe any decrease in the levels or activities of the cytosolic and nuclear proteins tested. This indicates that ISCU M141I is epistatic to FXN I154F (Figure 3 E and F).

### ISCU M141I prevents iron homeostasis dysregulation in the absence of FXN

Iron homeostasis is tightly linked to Fe-S clusters biogenesis, as iron is a key component of the clusters. Disruption of Fe-S cluster biogenesis often results in iron homeostasis dysregulation, characterized by mitochondrial iron accumulation and cytosolic iron depletion, a hallmark of Friedreich ataxia^37^. We therefore assessed whether expression of ISCU M141I in FXN-deficient cells could prevent iron dyshomeostasis.

We first evaluated activation of the iron regulatory proteins IRP1 and IRP2, which post-transcriptionally regulate iron homeostasis via binding to iron-responsive elements (IREs) in the mRNAs encoding proteins involved in iron trafficking and storage. IRP1 exists in an inactive form as cytosolic aconitase (ACO1) when containing a Fe_4_S_4_ cluster, and loss of the cluster converts it to its active IRE-binding form. IRP2 is stabilized under Fe-S cluster-deficiency conditions due to impaired degradation by the FBXL5 ubiquitin ligase, whose stability depends on cellular iron levels and whose E3 ligase activity requires a Fe_2_S_2_ cluster. IRE-binding activity of IRP1 and IRP2 was assessed by IRE pull-down followed by immunoblots. As expected, the FA line (*Iscu^+/+^; hFXN^I154F^*) showed increased binding of both IRPs, consistent with disrupted cytosolic Fe-S cluster availability (Figure 4A). Activation of the IRPs IRE-binding activities was absent in ISCU M141I expressing cells, regardless of FXN expression. Importantly, the inappropriate activation of IRP IRE-binding activity was normalized in the *Iscu^KI/KI^; hFXN^I154F^*line, indicating that ISCU M141I is sufficient to restore Fe-S cluster synthesis for the cytosol and prevent the aberrant activation of IRPs (Figure 4A).

**Figure 4:**
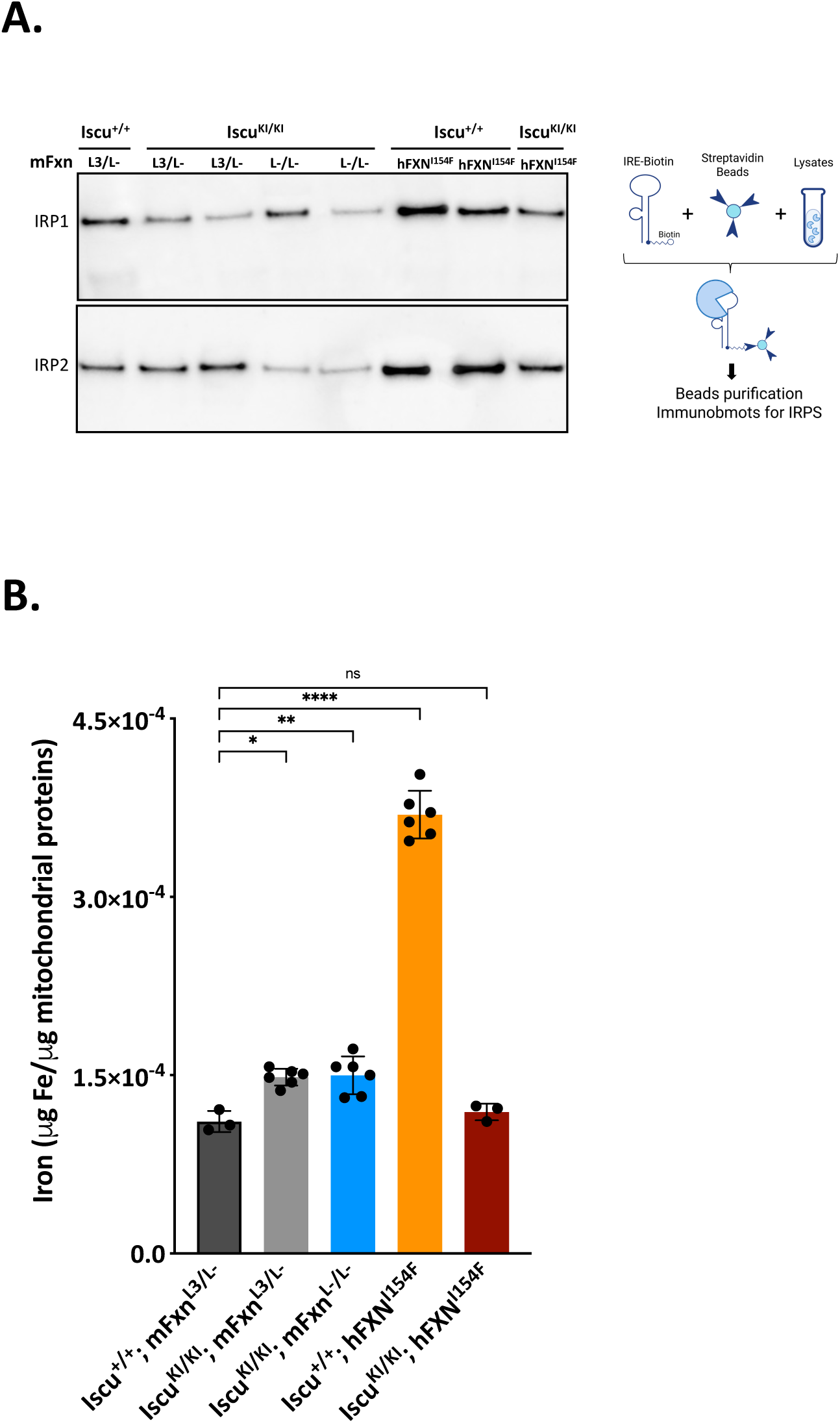
ISCU M141I prevents dysregulation of iron homeostasis in the absence of FXN. (A) Iron regulatory protein (IRP1 and IRP2) binding activities to Iron Responsive Element (IRE) was assessed using a biotin-streptavidin pull-down assay, allowing quantification of the amount of IRPs bound to a biotinylated ferritin-H IRE probe. (B) Mitochondrial iron content was measured by ICP-MS from control and different cell lines, as indicated. Iron levels were normalized to mitochondrial protein content to assess differences in mitochondrial iron accumulation. (n = 3 biological replicates, with two different clones for Iscu^KI/KI^; mFxn^L3/L-^ cells, Iscu^KI/KI^; mFxn^L-/L-^ cells bypassing lethality pooled together, and Iscu^+/+^; hFXN^I154F^ cells (“FA cells”)).

We then quantified mitochondrial iron content, normalized to mitochondrial protein levels. As previously reported^29^, the FA line showed a striking mitochondrial iron overload (Figure 4B). In contrast, mitochondrial iron levels in ISCU M141I expressing lines, including the FXN-deficient (*Iscu^KI/KI^; mFxn^L-/L-^*), remained close to normal despite a slight increase relative to control (Figure 4B). Interestingly, the *Iscu^KI/KI^; hFXN^I154F^* line also exhibited complete rescue of normal mitochondrial iron levels (Figure 4B). Together, these findings confirm that ISCU M141I restores iron homeostasis in FXN-deficient cells by preserving cytosolic Fe-S cluster biogenesis.

### ISCU M141I does not allow to bypass lethality due to absence of FXN in mice but delays the embryonic lethality

Having established that ISCU M141I variant can bypass lethality due to FXN deficiency *in vitro*, we next assessed whether this variant could also bypass embryonic lethality *in vivo*. Indeed, previous studies have shown that complete loss of FXN in mice leads to early embryonic lethality at E6.5 days^17^. We generated a transgenic mice line carrying the ISCU M141I mutation via CRISPR-Cas9 (Figure 5A). The modified allele (KI) harbors one G>C substitution in exon 5 of the Iscu gene, allowing the substitution of the methionine in position 141 into an isoleucine, along with two silent mutations creating a BstUI restriction site for genotyping (Figure 5A). Both heterozygous *Iscu^+/KI^* and homozygous *Iscu^KI/KI^*mice were viable, fertile and displayed no overt phenotypes. We then crossed this line with a line heterozygous for *mFxn* deletion (*mFxn^+/L-^*)^30^, generating the compound line *Iscu^KI/KI^; mFxn^+/L-^,* which were subsequently intercrossed to determine if the ISCU M141I variant allowed to bypass lethality (Figure 5A). According to Mendelian ratios, 25% of offsprings from *Iscu^KI/KI^; mFxn^+/L-^* intercross should be *Iscu^KI/KI^; mFxn^L-/L-^*. However, among > 30 litters genotyped postnatally, no *Iscu^KI/KI^; mFxn^L-/L-^* animals were recovered, indicating that ISCU M141I does not rescue embryonic lethality associated with FXN loss.

**Figure 5:**
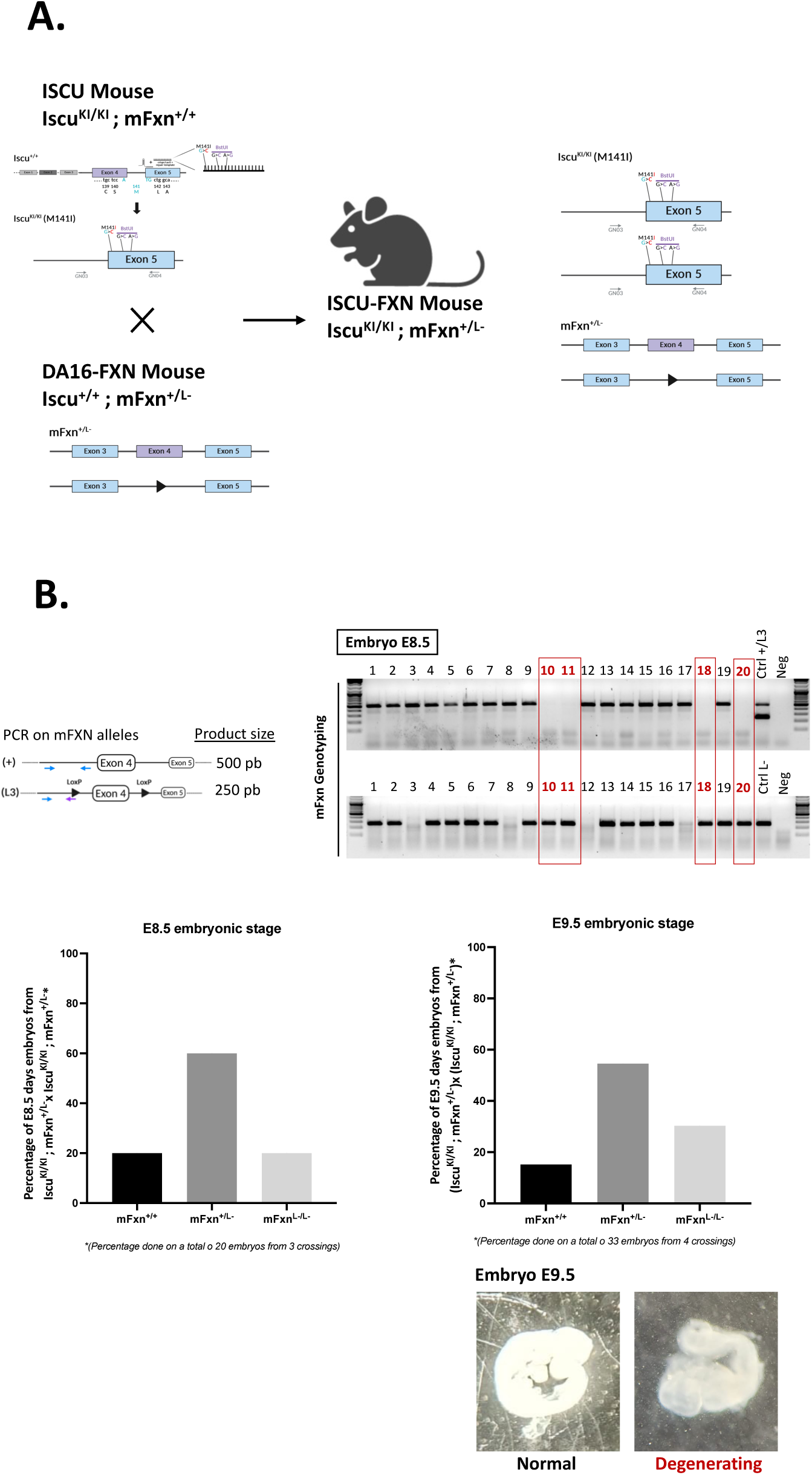
ISCU M141I does not rescue embryonic lethality caused by FXN deficiency in mice but delays its onset. (A) Schematic representation of mouse breeding strategies to generate the Iscu^KI/KI^; mFxn^L-/L-^ double mutant mouse model. (B) Genotyping example of mFxn alleles obtained from Iscu^KI/KI^; mFxn^+/L*-*^ intercross. Bar graphs show the distribution of genotype among embryos collected at embryonic day (E) 8.5 and E9.5. Representative images of morphologically normal and degenerating embryos at E9.5 stage are shown.

Given the beneficial effect of the ISCU M141I variant on cytosolic and nuclear Fe-S cluster distribution *in vitro*, we investigated whether this variant could extend the embryonic viability *in vivo* in the absence of FXN. To test this, we performed timed matings between *Iscu^KI/KI^; mFxn^+/L-^*mice and collected embryos at two developmental stages: E8.5 and E9.5. According to Mendelian ratio, we expected the following genotype distribution: 25% *Iscu^KI/KI^; mFxn^+/+^*, 50% *Iscu^KI/KI^; mFxn^+/L-^* and 25% *Iscu^KI/KI^; mFxn^L-/L-^*. At E8.5, no obvious morphological differences could be observed among the 20 embryos examined. Genotyping revealed that 20% were *Iscu^KI/KI^; mFxn^L-/L-^* (Figure 5B), indicating that the ISCU M141I variant permits survival of FXN-deficient embryos at least until this stage. At E9.5, we analyzed 33 embryos. Genotyping showed that 30.3% harbored the *Iscu^KI/KI^; mFxn^L-/L-^* genotype, consistent with Mendelian ratios. However, unlike at E8.5, embryos displayed a range of phenotypes, from healthy morphology to degenerating embryos and empty decidua (Figure 5B). None of the *Iscu^KI/KI^; mFxn^L-/L-^* embryos appeared healthy, indicating that despite the fact that the ISCU M141I delayed embryonic lethality due to FXN loss, it did not fully rescue viability.

These findings suggest that in the absence of FXN, the ISCU M141I variant extends embryonic survival by approximately 3 days, from E6.5 to E9.5, highlighting its partial compensation capacity *in vivo*.

## Discussion

In this study, we demonstrate that the ISCU M141I substitution, previously identified as a suppressor of frataxin deficiency in yeast, is sufficient to bypass the lethality caused by complete loss of FXN in dividing mammalian cells. This represents the first evidence that a single amino acid change in the scaffold protein ISCU can partially compensate for the essential role of FXN in mammalian Fe-S cluster biogenesis. In FXN-null murine fibroblasts, the ISCU M141I variant supported continued cell proliferation, despite reduced growth rates and persistent mitochondrial dysfunction. Importantly, these cells did not exhibit G2/M cell cycle arrest or DNA damage accumulation, indicating that cytosolic and nuclear Fe-S proteins involved in DNA metabolism remained functional. However, a broad deficit in mitochondrial Fe-S proteins was observed, consistent with the mitochondrial dysfunction.

Previous biochemical studies in yeast suggested that the methionine to isoleucine substitution in ISCU may enhance cysteine desulfurase activity^25^. Accordingly, the ISCU M141I variant could accelerate the rate of Fe-S cluster biogenesis in fibroblasts, which may be sufficient to restore activities of key Fe-S enzymes, particularly those required for cell division, in the absence of FXN. However, defects in mitochondrial Fe-S proteins persist, indicating that the ISCU M141I variant cannot fully compensate for the loss of FXN. Moreover, a major difference emerges when comparing the mammalian system to the yeast model. In the frataxin-deficient yeast expressing the Isu1 M141I variant, not only was growth restored, but full recovery of mitochondrial Fe-S proteins SDH and aconitase was also observed^25^. While cytosolic and nuclear Fe-S proteins were not investigated in the yeast study^25^, these findings suggest that the bypass mechanism conferred by the ISCU M141I variant operate differently in the unicellular yeast versus mammalian cells. One could hypothesize that the partial rescue of Fe-S proteins in mammalian cells lacking FXN depends on the number or the complexity of Fe-S clusters required by each protein. However, our data do not support this idea. For instance, mitochondrial proteins such as SDHB, which contains three Fe-S clusters of different stoichiometries (Fe_2_S_2_, Fe_3_S_4_, Fe_4_S_4_), and in FECH, which requires only a single Fe_2_S_2_ cluster, were comparably and severely affected. In contrast, cytosolic DPYD, which contains eight Fe_4_S_4_ clusters and nuclear POLD1, which requires a single Fe_4_S_4_ were fully functional. Thus, an increase rate of the Fe-S cluster biogenesis due to the ISCU M141I variant does not fully explain the compartment-selective rescue of Fe-S proteins in absence of frataxin. These findings may instead reflect that the ISCU M141I variant affects the distribution or prioritization of Fe-S clusters between mitochondrial and the cytosolic/nuclear targets.

Regulation of Fe-S cluster biogenesis, particularly the ISC machinery, remains poorly understood. Fe-S cluster biogenesis is closely linked to cellular metabolism, iron homeostasis and ROS production. Deficiency in Fe-S cluster synthesis is associated with metabolic disorders^38^, and increased Fe-S turnover is seen in highly proliferative cancer cells under oxidative stress^39,40^. Reduced Fe-S cluster production, as in Friedreich’s ataxia, leads to iron accumulation^37^. ISCU is regulated by different metabolic signals: upon DNA damage, p53 enhances its expression to control iron homeostasis^41^, miR-210 represses it under hypoxia^42^, and mTORC1 stabilizes it via phosphorylation^43^. However, we found no difference in ISCU levels between cells carrying wild-type and M141I variant without FXN, yet the M141I variant prevented mitochondrial iron accumulation, likely through its ability to restore Fe-S cluster incorporation into nucleocytoplasmic proteins, including IRP1 and FBXL5. Overall, our results suggest that the thus far reported regulatory mechanisms are not involved and do not explain the compartment-specific effects observed.

Our results raise the intriguing possibility of compartment-specific regulation of Fe-S cluster allocation, a process that remains poorly explored and understood. While mitochondrial and cytosolic Fe-S protein maturation both depend on the mitochondrial ISC machinery, the export of the sulfur/iron containing intermediates for cytosolic/nuclear Fe-S protein biogenesis via the CIA pathway introduces a layer of complexity. Our data show that the ISCU M141I variant restores early CIA components (such as GLRX3), support ^55^Fe incorporation into cytosolic targets, and prevents IRP activation and mitochondrial iron overload, all phenotypes that are hallmarks of impaired cytosolic Fe-S cluster assembly. These findings suggest that the ISCU M141I may enhance the export or delivery of Fe-S cluster precursors to the CIA machinery.

The structural context of the M141 residue offers potential mechanistic insights. This methionine on ISCU lies adjacent to the conserved LPPVK motif, which mediates the interaction with the HSPA9/HSC20 chaperone complex required for downstream Fe-S cluster transfer^44^. Cryo-electron microscopy studies have shown that FXN binding to the ISC complex promotes displacement of ISCU and conformational changes in the LPPVK loop^13^. Substituting the methionine with isoleucine could potentially affect the interaction of ISCU with the chaperone complex via the LPPVK motif, altering its binding dynamics and perhaps favoring the export of (Fe-S)_int_. Supporting this hypothesis, *in vitro* studies involving the human ISCU M140I variant (the equivalent mutation) show that the methionine into isoleucine substitution increases the rate of Fe-S cluster transfer to the mitochondrial glutaredoxin 5, GLRX5^28^, a key protein bridging the ISC and CIA pathways. Indeed, a recent study in yeast identified the central role of GLRX5 in Fe-S cluster trafficking, demonstrating its necessity for both mitochondrial and cytosolic/nuclear Fe-S proteins^45^. While biochemical studies of the ISC machinery have provided valuable insights into the ISC complex dynamics during de novo assembly of Fe-S clusters, the specific interactions between Fe-S-loaded ISCU, the chaperone complex, and GLRX5 remain largely undefined. In particular, it will be interesting to determine whether the methionine to isoleucine substitution alters these interactions, potentially biasing (Fe-S)_int_ export towards the cytosol.

Importantly, the ISCU M141I variant was unable to overcome the embryonic lethality of FXN deficiency *in vivo*, despite significantly delaying it from E6.5 to E9.5. We can hypothesize that restoration of cytosolic and nuclear Fe-S proteins, particularly those required for cell proliferation, combined with functional maternal mitochondria, supports initial development. However, by E8.5-E9.5, when the embryo becomes largely dependent on its own mitochondria, persistent mitochondrial dysfunction likely leads to embryonic development arrest.

Altogether, our results reveal a previously unrecognized compartment-specific rescue of Fe-S cluster-dependent processes by the ISCU M141I variant in mammalian cells, raising for the first time the possibility of compartmental regulation of Fe-S cluster biogenesis. This prompts key questions: How does the cell prioritize Fe-S cluster allocation between cytosolic/nuclear and mitochondrial compartments? Which molecular players mediate this regulation? Could these mechanisms differ in post-mitotic cells, those primarily affected in FA? The ISCU M141I variant provides a valuable tool to investigate these questions and to dissect the molecular determinants governing the distribution and regulation of Fe-S clusters across cellular compartments.

## Limitations of the study

A first limitation of our study is that the differential rescue of mitochondrial versus cytosolic/nuclear Fe-S proteins by the ISCU M141I variant in the absence of FXN was demonstrated only in one cellular model, namely immortalized fibroblasts. Additional investigations in other cell types will be required to validate and generalize the hypothesis of compartment-specific regulation of Fe-S cluster biogenesis. A second limitation lies in the intrinsic constraints of using mitotic cells. In this setting, it is not possible to generated complete FXN knockout controls carrying wild-type ISCU, given the well-established lethality associated with the absence of frataxin. Finally, our study is limited by the current absence of structural and biochemical characterization of the ISCU M141I variant, both as an isolated protein and in complex with is partners within the Fe-S clusters biogenesis machinery. Such studies will be essential to elucidate the precise molecular basis of its functional properties.

## Acknowledgement

This work was supported by the Agence Nationale pour la Recherche (ANR# 17-CE12-0033) (to H.P.) and by the Intramural Research Program of the *Eunice Kennedy Shriver* National Institute of Child Health and Human Development (to N.M.). The ISCU M141I variant mouse was generated at PHENOMIN-ICS by a grant from PHENOMIN (“programme Investissements d’Avenir” (ANR-10-INBS-07 PHENOMIN)).

## Authors Contributions

V.M. conceived and conducted most of the experiments, analyzed the data, prepared the figures and wrote the manuscript. N.M. performed all the experiments for the analysis of nuclear and cytosolic Fe-S proteins and for the mitochondrial iron content measurement, edited the manuscript and provide critical input; N.D. generated the CRISPR-Cas9 engineered murine fibroblast (Iscu^+/+^; mFxn^L3/L^) and preliminary data of the study; A.H. assisted with the experimental studies; L.D. performed dissection of embryos at E8.5 and E9.5; L.R. performed dissection of embryos at E8.5 and E9.5; L.M. performed mouse strains maintenance and crossings; M.C.B. designed the CRISPR-Cas9 mutagenesis protocol for the ISCU mouse generation; A.E. initiate the generation of the CRISPR-Cas9 engineered murine fibroblast (Iscu^+/+^; mFxn^L3/L^). A.M. initiate and designed the CRISPR-Cas9 mutagenesis protocol for the murine fibroblasts; H.P. conceived and supervised the research, obtained funding and wrote the manuscript.

---

**Supplementary Data 1:**
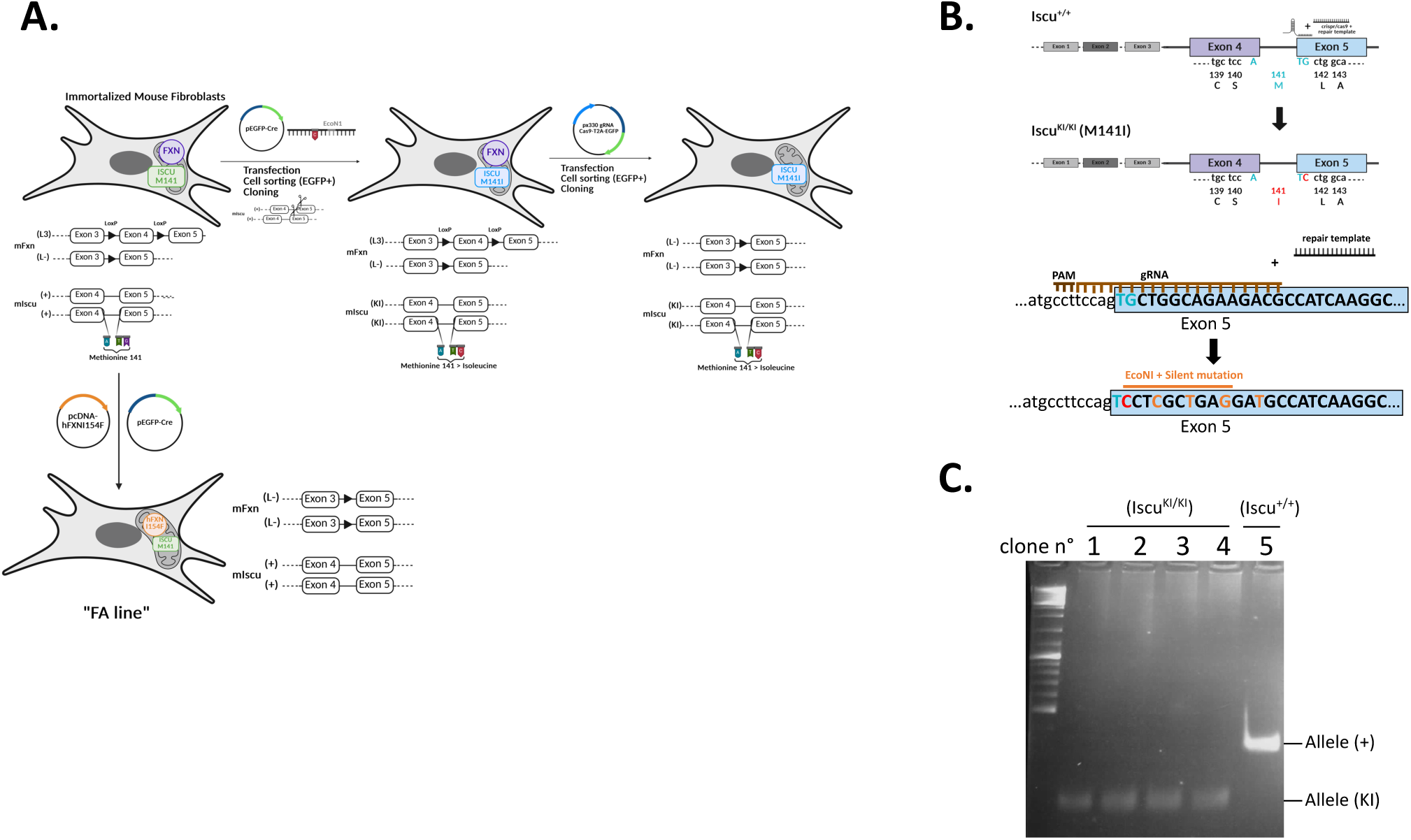
Insertion of *Iscu* G to C substitution (M141I) by CRISPR-Cas9 in immortalized mouse fibroblasts. (A) Schematic overview of the different cell lines generated in the study from immortalized mouse fibroblasts carrying a conditional allele on the Fxn gene (L3/L-). The G>C substitution in the Iscu gene (resulting in the M141I amino acid change) was introduced using CRISPR-Cas9 combined with a homology-directed repair (HDR) template. Deletion of exon 4 of the murine frataxin (mFxn) gene was achieved by Cre-recombinase-mediated excision. (B) Detailed schematic of the CRISPR-Cas9 strategy used to introduce the G>C substitution in exon 5 of *Iscu*. (C) Screening of the edited clones. PCR amplification of a 337 pb product surrounding exon 5 of the Iscu gene was followed by EcoNI restriction digestion. The presence of efficient recombination with the HDR template, and therefore the G> C substitution, is identified by the presence of two EcoNI digestion products (173 and 167 pb, indistinguishable from each other on gel).

**Supplementary Data 2:**
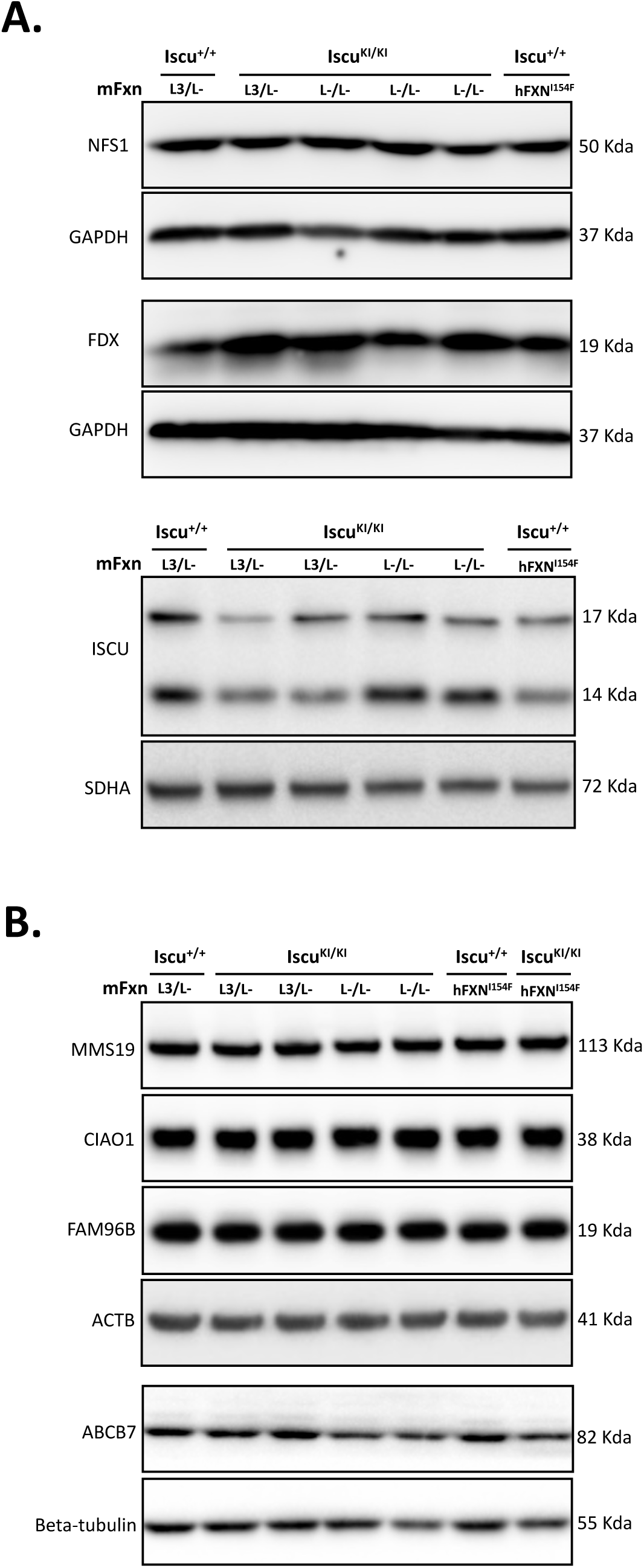
Western blot analysis of components of the ISC and CIA machineries. (A) Western blot analysis of the mitochondrial ISC machinery members NFS1, FDX and ISCU in the different cell lines. (B) Western blot analysis of the CIA machinery members MMS19, CIAO1, FAM96B and ABCB7 in the different cell lines.

**Supplementary Data 3:**
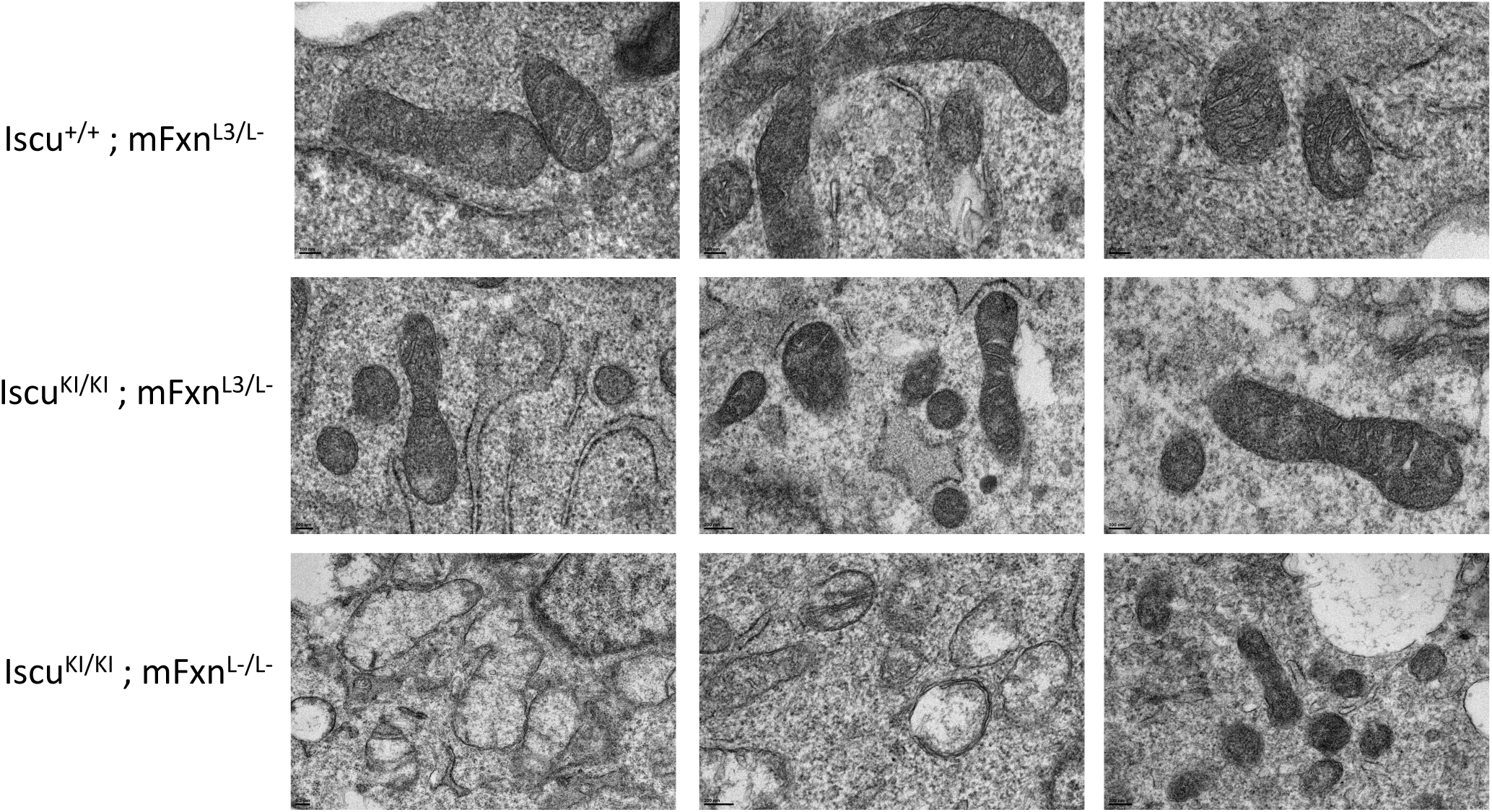
Electron microscopy analysis. Representative pictures of mitochondrial morphology and health by electron microscopy in Iscu^+/+^; mFxn^L3/L-^ control cells, Iscu^KI/KI^; mFxn^L3/L-^ cells, and Iscu^KI/KI^; mFxn^L-/L-^ cells bypassing lethality due to FXN loss.

